# INC-Seq: Accurate single molecule reads using nanopore sequencing

**DOI:** 10.1101/038042

**Authors:** Chenhao Li, Kern Rei Chng, Esther Jia Hui Boey, Amanda Hui Qi Ng, Andreas Wilm, Niranjan Nagarajan

## Abstract

Nanopore sequencing provides a rapid, cheap and portable real-time sequencing platform with the potential to revolutionize genomics. Several applications, including RNA-seq, haplotype sequencing and 16S sequencing, are however limited by its relatively high single read error rate (>10%). We present INC-Seq (Intramolecular-ligated Nanopore Consensus Sequencing) as a strategy for obtaining long and accurate nanopore reads starting with low input DNA. Applying INC-Seq for 16S rRNA based bacterial profiling generated full-length amplicon sequences with median accuracy >97%. INC-Seq reads enable accurate species-level classification, identification of species at 0.1% abundance and robust quantification of relative abundances, providing a cheap and effective approach for pathogen detection and microbiome profiling on the MinION system.

## 1. Introduction

The MinION sequencing platform introduced by Oxford Nanopore Technologies Inc. (ONT) is a portable and inexpensive device that generates long reads in real time, and can thus further democratize genomics [1]. Compared to short accurate reads on second-generation sequencing platforms (e.g. Illumina, Ion Torrent), nanopore sequencing shares the feature of other third-generation platforms in that it trades off base level accuracy for much longer reads. However, several genomics applications ideally require both long and highly accurate reads (e.g. RNA-seq [2], haplotype sequencing [3] and 16S sequencing [4]) to help discriminate between very similar DNA sequences.

Raw single-strand reads (1D) on the MinION platform have been reported to have error rate >20%, but typically more accurate reads (10-15% median error rate) are generated by a protocol to read the complementary strand as well to compute an *in silico* consensus (2D reads) [5]. Extending this idea further, experimental protocols have been developed for other sequencing platforms to boost read accuracy by repeatedly sequencing the same parent template molecule, and obtaining more reliable consensus reads [6–9]. For e.g., BAsE-Seq [6] uses a sophisticated library preparation protocol to tag each template with a unique barcode, resolving short barcodes with one end of a paired-end Illumina read, and using the other end to reconstruct a consensus template read. PacBio circular consensus sequencing [7] (also known as SMRT CCS) reads through circularized templates repeatedly by synthesis, but cannot be directly applied to nanopore sequencing which requires a single stranded linear template to be threaded through the nanopore channel. Circ-Seq [8, 9] generates linear templates (3 repeating units on average) with a protocol that is customized for short templates to be sequenced on the Illumina short-read platform.

We introduce INC-Seq (for Intramolecular-ligated Nanpore Consensus Sequencing) that employs rolling circle amplification (RCA) of circularized templates to generate linear products (with tandem copies of the template) that can then be sequenced on the nanopore platform. Circularization, amplification and nanopore sequencing have several challenges including (i) generation of hybrid molecules (chimeras), (ii) introduction of polymerase errors, (iii) amplification bias from exponential amplification, and (iv) input DNA requirements of the nanopore platform. These issues are addressed in INC-Seq’s experimental protocol, allowing for the construction of consensus single molecule reads with median accuracy >97% using low input DNA. Applying INC-Seq to rRNA based community profiling allowed for accurate classification at the species level and identification of rare species in the community. INC-Seq profiles were found to be robust and well correlated with known abundances. As 16S rRNA sequencing is often the method of choice when working with samples with high contamination, INC-Seq provides a cost-effective approach for pathogen detection and microbiome profiling. In addition, INC-Seq provides an avenue for the use of nanopore sequencing in many genomic applications that require high sequence accuracy.

## 2. Materials and Methods

### 2.1. Bacterial DNA extraction, PCR amplification of 16S rRNA and construction of synthetic communities

Bacterial DNA was obtained from DSMZ Germany for *Parvimonas micra* (DSM 20468), *Streptococcus cristatus* (DSM 8249), *Streptococcus oralis* (DSM 20627), *Faecalibacterium prausnitzii* (DSM 17677), *Neisseria subflava* (DSM 17610), *Bifidobacterium longum* subsp. *longum* (DSM 20219), *Fusobacterium nucleatum* subsp. *polymorphum* (DSM 20482) and *Helicobacter pylori* (DSM 21031). Bacterial cultures were obtained from ATCC for *Staphylococcus epidermidis* (ATCC 12228), *Salmonella typhimurium* subsp. *enterica* (ATCC 13311), *Klebsiella pneumoniae* subsp. *pneumoniae* (ATCC 700721), *Streptococcus pyogenes* (ATCC 49117) and *Staphylococcus aureus* subsp. *aureus* (ATCC 31240). Bacterial cultures were grown in ATCC recommended media and DNA was extracted using a genomic DNA isolation kit (Bioline, bio-52066), according to manufacturer’s protocol. PCR amplification of 16S rRNA was carried out using previously designed primers [10], 338F (5’ Phos –ACTYCTACGGRAGGCWGC– 3’) and 1061R (5’ Phos –CRRCACGAGCTGACGAC– 3’), using Phusion High-Fidelity PCR Master Mix (F531, Thermo Fisher Scientific) according to the manufacturer’s instructions. The conditions for PCR were set up as follows: Initial denaturation at 98 °C for 30s, 15 cycles of 98 °C for 10s with annealing at 59 °C for 30 s and extension at 72 °C for 30s, and final extension at 72 °C for 5 min. PCR products were purified with Agencourt AMPure XP beads (Beckman Coulter). Two synthetic bacterial communities were constructed by pooling (by mass) 1ng of 16S rRNA DNA in total according to defined relative abundances (**Table 1** and **Table 2**). The “simple” community consists of three bacteria while the “ladder” community consists of 10 bacteria with large variation in relative abundances.

**Table 1.** INC-Seq estimated abundances for a simple synthetic community

| Species | Reference (GenBank ID) | Relative abundance |  |  |
| --- | --- | --- | --- | --- |
|  |  | Defined | SMRT | ONT 2D |
| *Streptococcus oralis* | KT932114.1 | 0.6 | 0.55 | 0.64 |
| *Streptococcus cristatus* | KF933778.1 | 0.3 | 0.39 | 0.31 |
| *Parvimonas micra* | KP944178.1 | 0.1 | 0.06 | 0.05 |

**Table 2.** Species level profiling for a “ladder” synthetic community

| Species | Relative abundances |  |
| --- | --- | --- |
|  | Defined | INC-Seq |
| *Staphylococcus aureus* | 0.3620 | 0.6310 |
| *Streptococcus pyogenes* | 0.3200 | 0.1476 |
| *Helicobacter pylori* | 0.1600 | 0.0959 |
| *Fusobacterium nucleatum* | 0.0800 | 0.0369 |
| *Bifidobacterium longum* | 0.0400 | 0.0424 |
| *Klebsiella pneumoniae* | 0.0200 | 0.0092 |
| *Salmonella typhimurium* | 0.0100 | 0.0221 |
| *Staphylococcus epidermidis* | 0.0050 | 0.0074 |
| *Neisseria subflava* | 0.0020 | 0.0037 |
| *Faecalibacterium prausnitzii* | 0.0010 | 0.0018 |
| *Klebsiella variicola* */ *pneumoniae* | 0 | 0.0018 |

\* *Variicola* and *pneumoniae* differ by one mismatch.

### 2.2. Circular ligation and rolling circle amplification for construction of INC-Seq libraries

Circular ligation of pooled (by mass) DNA was performed in 3 mL reaction volume with T4 DNA ligase (M0202, New England Biolabs) according to manufacturer’s protocol. The mixture was incubated at 16 °C for 6 h before heat inactivation of T4 DNA ligase at 65 °C for 10 min. Ligated products were purified and concentrated using AMPure XP beads. Plasmid-Safe ATP-Dependent DNase (E3101, Epicentre) digest was performed according to manufacturer’s protocol at 37 °C for 1 h to remove non-circularized products. DNase was then heat inactivated at 70 °C for 30 min. Products were purified and concentrated using AMPure XP beads. The circular DNA template molecules were added to phi29 DNA polymerase reaction buffer (B0269, New England Biolabs) with 0.25 μg/μL BSA, 5 μM exo-resistant random primers (S0181, Thermo Fisher Scientific) and 10 U phi29 polymerase (M0269, New England Biolabs) in a total volume of 20 μL. The conditions for rolling circle amplification (RCA) were set up as follows: 95 °C for 3 min, 30 °C (ramp rate of 0.1°C/s) for 6 h and 65 °C for 10 min. The time taken for amplification was optimised to yield substantial RCA products, while minimising the possibility of sequences produced by unspecific priming of random primers. RCA products were finally purified with 0.45x volume of AMPure XP beads.

### 2.3. PacBio library preparation and sequencing

The RCA product of the simple community was purified and concentrated with 1x volume AMPure^®^ PB Beads (100-265-900, Pacific Biosciences) giving 7 μg of purified product, determined by Qubit 2.0 fluorometer (Life Technologies). The purified product was gently sheared (to reduce hyperbranched DNA structures) using Covaris^®^ g-TUBE™ device (Covaris) at 6000 rpm for 1.5 min. To verify the size of fragments after shearing, sheared and un-sheared products were run on a 0.5% (w/v) agarose gel. The sheared RCA product was purified with 0.45x AMPure^®^ Beads and was used to prepare the 10 kb SMRTbell™ template in accordance with the Pacific Biosciences procedure for ‘10 kb Template Preparation and Sequencing’. BluePippin™ (Sage Science) was used to size-select libraries of sizes ranging from 6-10 kb. The distribution of library sizes was first assessed on Agilent 2100 Bioanalyzer (Agilent Technologies, Inc.) with the DNA 12000 kit before they were sequenced on a PacBio sequencer, using P5-C3 chemistry coupled with MagBead Standard Seq v2 collection protocol.

### 2.4. Nanopore library preparation and MinION sequencing

RCA DNA products were fragmented using Covaris^®^ g-TUBE™ (Covaris) centrifuged at 3200 rpm × g for 3 min. The Oxford Nanopore Technologies SQK-MAP006 protocol was followed, with some modifications to the Run B protocol from Urban *et al.* [11]. Briefly, starting material of 8 μg was used, the RCA products were handled using wide-bore tips and 0.4x AMPure XP beads purification was performed after DNA repair and end prep. As more starting material was used, DNA repair, end prep and ligation were performed in double the volume with double the reagents. All elutions were carried out for 20 min at 37 °C. Samples were processed with SQK-MAP006 kits and run on SQK-MAP006 flow cells on a MinION MK1 device. The raw fast5 reads were uploaded to the online server for basecalling through the Metrichor software (version 1.48). 2D passed reads in fasta format were extracted using poretools (version 0.5.1) [12].

### 2.5. Anchor-based consensus construction for INC-Seq reads

For each original INC-Seq read, subsequences were extracted using non-overlapping sliding windows of 500 base pairs (till half the read length) to serve as “anchors” (i.e. subsequences 1-500, 501-1000, 1001-1500…). The anchors were aligned to the read with BLASTN (for high sensitivity [13]). Alignments covering less than 80% of an anchor were discarded. Anchors with the most number of alignments were used to extract repeating segments in the read. The starting positions of two adjacent anchors define a candidate repeating segment. To extract segments that are as complete as possible, the segment end corresponding to the 5’-end anchor was extended according to the clipped bases of the anchor at its 5’-end. For each read, the longest run of consecutive segments with the same orientation and similar lengths (up to 5% difference from the median length) was used to construct a consensus sequence. Each segment extracted from a run was tested as a potential backbone by aligning other segments against it with BLASTN. The segment with the most number of alignments (breaking ties using the average percentage identity of alignments) was used as the final backbone. Reads having less than six aligned segments were discarded. BLASTN alignments were converted to BLASR’s m5 format and a consensus sequence constructed using PBDAGCON (-t 2 -c 1 -m 5; git commit #de1cf85; https://github.com/PacificBiosciences/pbdagcon).

### 2.6. Species profiling using simulated reads

A customized 16S reference database was constructed by trimming sequences in the SILVA database (release 123) [14] to the amplicon regions (i.e. from V3 to V6) [10] and clustering trimmed sequences at 97% identity using vsearch. (version 1.9.3; https://github.com/torognes/vsearch). Ten representative OTU sequences were selected at random (**Supplementary File 1**) from the customized database and ONT 2D reads were simulated from each OTU sequence according to the error profile reported by Camilla *et al.* [5] (3% mismatch, 4% insertion, 5% deletion) using PBSIM (--length-mean <reference_16S_length> --length-sd 0) [15]. To simulate RCA reads, synthetic amplified sequences were created by concatenating each reference sequence 40 times. Simulated reads were generated using PBSIM (--length-mean <read_length> --length-sd 0) with the same error profile, where “read_length” was determined by the number of copies required for correction. Simulated reads were mapped to the correct reference and the customized database to calculate percentage identity and classification accuracy respectively.

### 2.7. Estimating chimera rates

In INC-Seq, chimeras can result from (i) inter-molecular ligation of different 16S sequences and (ii) template switching during amplification. To estimate respective chimera rates, reference 16S sequences for *Streptococcus cristatus* and *Parvimonas micra* were trimmed to the amplicon region and then aligned to PacBio and nanopore reads from the simple community using BLASTN (version 2.2.28+) [16]. Alignments with the lowest E-value were selected for each region of the reads, and alignments covering less than 80% of the 16S query were discarded. As *Streptococcus oralis* has 97% identity to *Streptococcus cristatus*, we grouped the two *Streptococcus* species in this analysis.

For inter-molecular ligation, reads containing alternating patterns of both references (at least 2 copies of each) were considered as evidence and the intermolecular ligation rate was estimated to be:

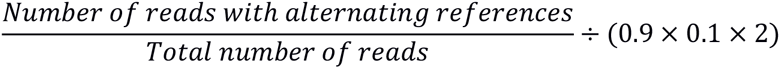

The additional term (0.9 × 0.1 × 2) accounts for the fact that we only observe inter-molecular ligations across genera. Similarly, for template switching, the number of reads containing both references but without an alternating pattern were considered as evidence and the template switching rate was estimated to be:

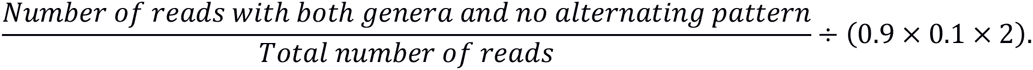

### 2.8. Assessing the quality of corrected sequences

The simple synthetic community was used to estimate the error rates of raw and corrected reads. For raw reads, the three trimmed reference 16S sequences (*S. oralis, S. cristatus* and *P. micra*) were aligned against each read with BLASTN and for each region alignments with the lowest *E*-value were selected. Error rate estimates (i.e. the percentage of mismatches, insertions and deletions) for each read were calculated by the ratio between the total number of errors and the alignment length. For corrected reads, each read was aligned against the three trimmed references and only the best alignment by *E*-value was kept. The error rate estimates were calculated as the ratio between the total number of errors and the total alignment length.

### 2.9. Species level profiling with INC-Seq corrected reads

INC-Seq corrected reads were self-concatenated (to recover the correct starting and end points) and mapped against the customized SILVA database (see **Section 2.6**) with BLASTN (the database was restricted to sequences annotated at species level and de-duplicated with vsearch). Alignments were filtered by reference coverage (>98%) and percentage identity (>98%). Species names were assigned according to the hit with highest identity to the query. The abundance of each reference species was estimated as the proportion of corrected reads that were mapped to that reference.

## 3. Results

### 3.1. The INC-Seq workflow

The INC-Seq workflow is based on template circularization, rolling circle amplification, nanopore sequencing and *in silico* consensus construction (**Figure 1**). The experimental procedure for generating INC-Seq libraries is summarized in **Fig. 1A**. Briefly, ligation conditions were optimized for intra-molecular ligation of long DNA molecules to generate predominantly self-ligated circular DNA templates. Non-circular DNA molecules are removed from the ligation mixture by exo-nuclease treatment. Following this, rolling circle amplification is carried out to produce long DNA molecules comprising of multiple repeating units. These long DNA products are then gently sheared and prepared for nanopore sequencing.

**Figure 1.**
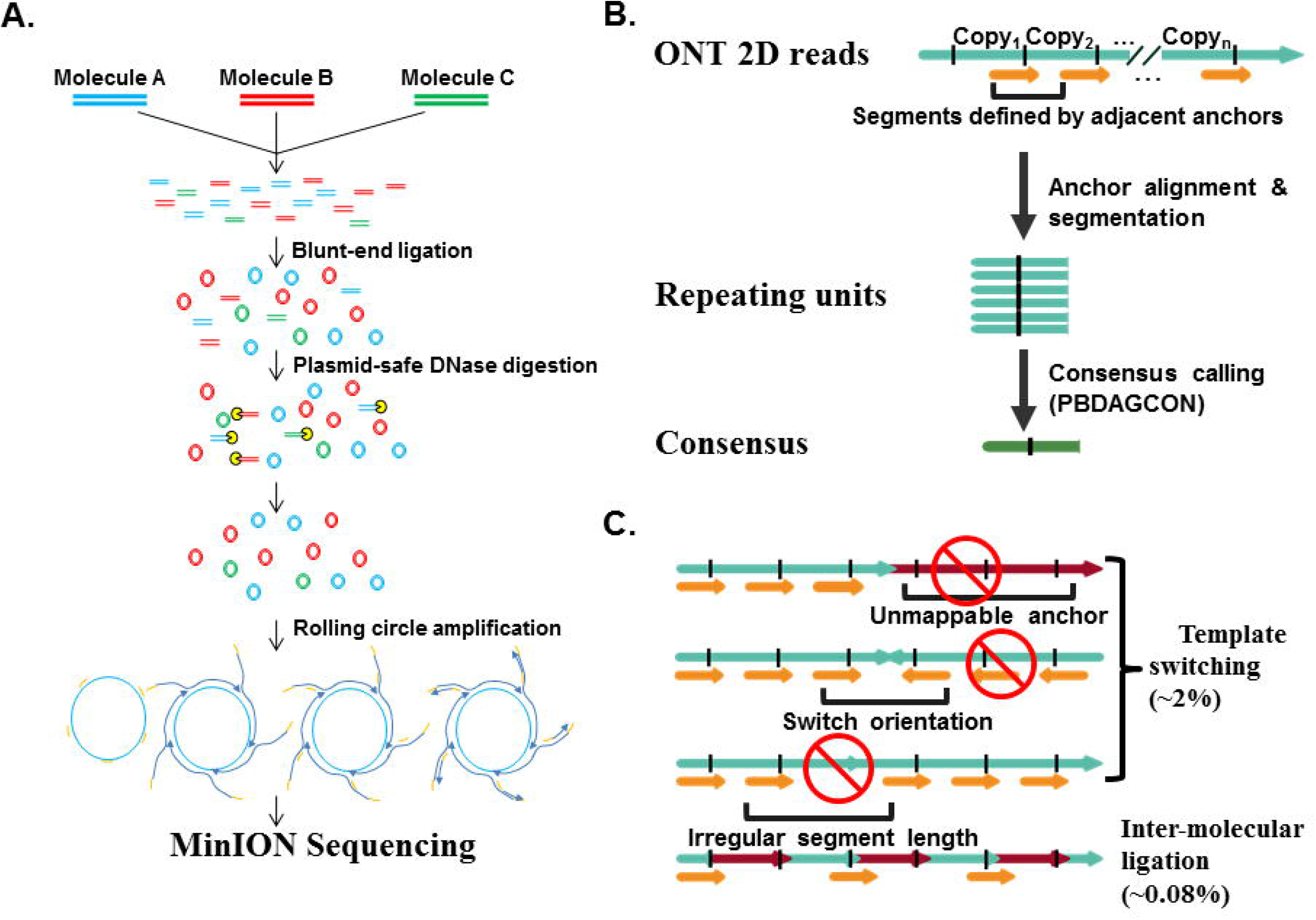
Overview of the INC-Seq workflow. (A) Template molecules are circularized in optimized conditions and the remaining linear molecules are removed. The circular products are amplified with RCA and sequenced on the MinION platform. (B) A subsequence from the raw nanopore reads is used as an anchor to scan the entire read for the location of repeating units. The repeating units flanked by adjacent anchor starting points are aligned and a consensus sequence is constructed. (C) In INC-Seq library preparation, chimeras are expected through inter-molecular ligation and template switching. Chimeras from template switching are likely to be detected by the anchor mapping protocol. Chimeras from inter-molecular ligation were observed to be rare under the experimental conditions used in INC-Seq.

To correct raw ONT reads, we designed a bioinformatics pipeline that extracts repeating sequence segments and corrects them by constructing a consensus sequence (see **Section 2.5** for details; **Figure 1B**). For each long read, an “anchor” subsequence of 500bp is selected and aligned to the read. Regions between two adjacent anchors then define the repeating segments. The repeating segments are then aligned to construct a consensus INC-Seq read.

The correction protocol in INC-Seq requires that each read consists of tandem repeats of the same template, and the presence of chimeric repeats can lead to a poor or chimeric consensus. Chimeras can be formed in the INC-Seq protocol due to inter-molecular ligation (during circularization) or due to template switching (during RCA). Template switching results in discordant mapping of an anchor (i.e. unmappable anchor, orientation switch or irregular anchor distance; **Figure 1C**). Chimeras from template switching can thus be detected from the concordance of mapped anchors, except in the case where switching happens between similar templates while preserving orientation and segment length. Chimeras from inter-molecular ligation are not directly discovered by the pipeline but will generate consensus sequences that are more than twice as long as expected. Chimeras from inter-molecular ligation were also observed to be rare under the experimental conditions used in INC-Seq (see **Section 3.3**) and can be reduced further by carrying out ligations in more dilute conditions (**Figure 1C**).

### 3.2. INC-Seq can reduce sequencing errors and significantly improve 16S OTU classification

INC-Seq was applied to 16S rRNA profiling as a proof-of-concept, due to its widespread use for bacterial identification in complex communities and samples with high non-microbial contamination. 16S sequences can be highly similar and thus both long and accurate rates are ideal for specific taxonomic classification. To evaluate if raw ONT 2D reads can be used for accurate species classification, we simulated reads from 10 OTUs and evaluated accuracy of classification (**Figure 2**). Classification accuracy was found to be highly variable (48-100%) with a median accuracy of 81% at the species level, and with many reads mapping to OTUs from a different taxonomic order (**Supplementary File 1**). Note that this is an idealized setting with the correct reference in the database.

**Figure 2.**
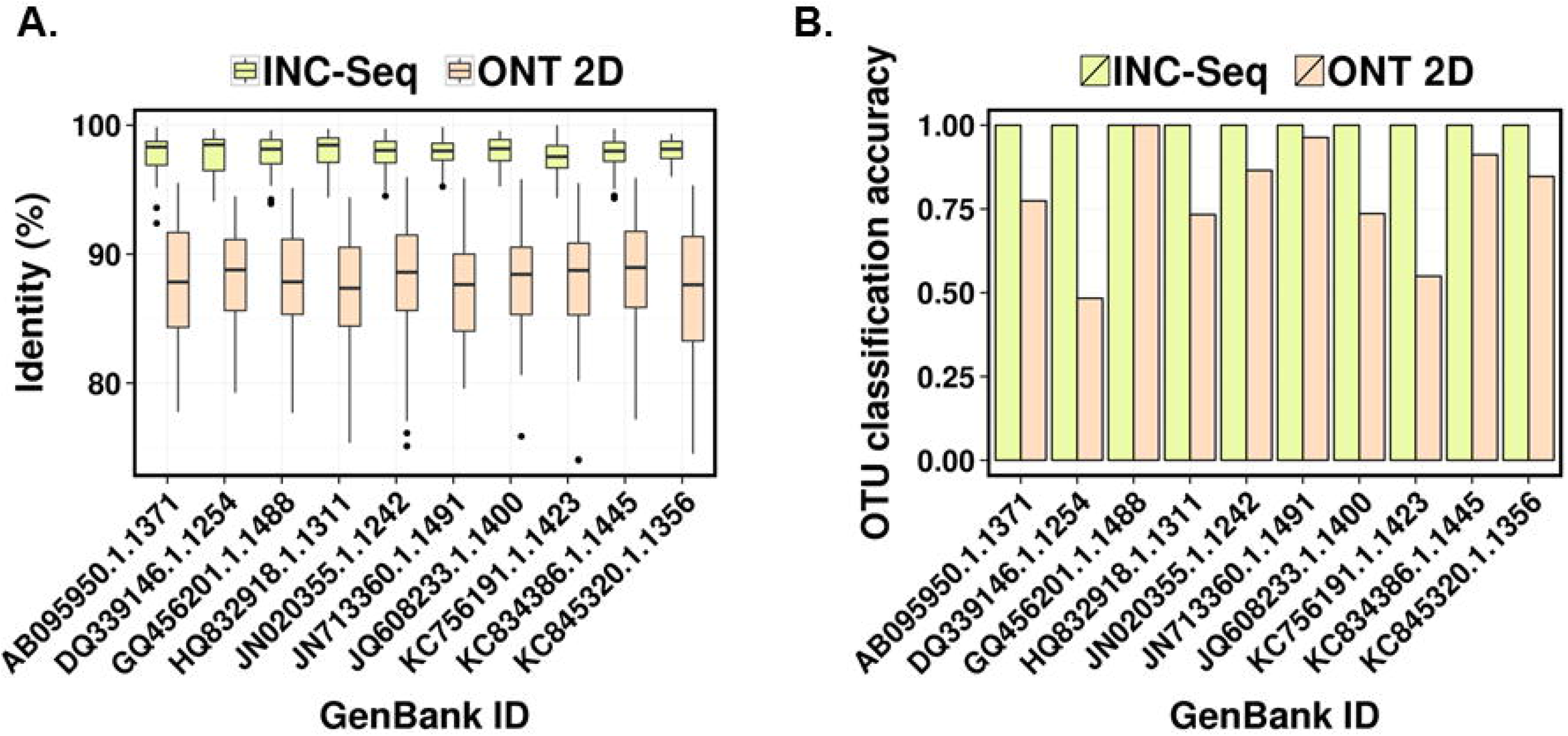
INC-Seq can significantly reduce read errors and improve OTU classification accuracy. Ten representative OTUs were selected from a customized database. Synthetic reads were generated for each reference OTU as well as artificial RCA products (i.e. INC-Seq libraries). (A) INC-Seq can reduce read errors 10-fold compared to raw ONT 2D reads. (B) Raw ONT reads frequently map to an incorrect reference while INC-Seq reads are more often assigned to the correct reference.

In comparison, simulated INC-Seq reads with various lengths, which were corrected in a reference free manner, reduced sequencing errors and improved 16S OTU classification accuracy (**Supplementary Figure 1**). INC-Seq reads containing six segments could be corrected to 98% median accuracy (average across 10 reference OTU sequences), reducing read error by a factor of six (**Figure 2A**). On average 82% of reads could be corrected in this fashion (**Supplementary File 1**), and all corrected reads could be perfectly assigned to the correct reference OTU (**Figure 2B**). Therefore, these values were used as thresholds (6 copies for correction and 98% for species level classification) for the synthetic communities below.

### 3.3. Assessing chimera rates in INC-Seq libraries with PacBio sequencing

PacBio SMRT sequencing of an INC-Seq 16S rRNA library (hyper-variable regions V3 to V6) for a simple synthetic bacterial community (i.e. *S. cristatus, S. oralis* and *P. micra*; **Table 1**) was used to estimate chimera rates for INC-Seq and to further evaluate the practicality of this protocol. Raw PacBio reads have a lower mismatch error rate (∼1%) [17] and can thus facilitate the identification of chimeric constructs. In addition, the median read lengths obtained here were slightly longer than those for MinION reads (4.7 vs 3.6 kbp; even though much longer reads can be obtained on the MinION system; **Supplementary Figure 2**) allowing for slightly better chimeric read analysis.

By mapping known, distinct reference 16S rRNA amplicon sequences to SMRT reads (see **Section 2.7**) the inter-molecular ligation rate was estimated to be 0.06% (**Supplementary Table 1**). In addition, no consensus INC-Seq reads that were longer than expected (>1 kb) were found to match the 16S rRNA gene, suggesting that the requirement of having 6 segment copies (and therefore raw read length >9 kb) may have filtered out chimeras from inter-molecular ligation.

A majority of chimeric constructs were estimated to come from template switching with up to 3% of reads in the SMRT library being attributed to it (**Supplementary Table 1**). Due to the conservative consensus construction approach used in INC-Seq (**Figure 1C**), such chimeras are expected to largely limit the effective length of runs of consecutive segments that are analysed (see **Section 2.5**), and thus the fraction of raw reads for which consensus reads are generated (as ≥6 segments are required for accurate consensus reads; **Supplementary Table 1**). As read lengths improve, this bottleneck is likely to be alleviated. Consensus reads from SMRT INC-Seq were found to cover the full length amplicon (median: 99%; mean: 98%) and improved median identity of SMRT reads from 84% to 98% (**Supplementary Figure 3A, B**). Mismatch, insertion and deletion error rates were reduced significantly (one-sided wilcoxon test *p*-value < 10^-16^; **Supplementary Figure 3C)**, and the median total error rate was reduced to below 1% when more than 15 segment copies were used to construct a consensus (**Supplementary Figure 3D**). The proportion of reads with significantly lower identity than average (2 standard deviations away at 92%) was low (2.6%), indicating that few INC-Seq reads are impacted by uniformly low accuracy or errors associated with chimeric sequences. Note that consensus sequences here were constructed from called bases and therefore accuracy is likely to improve further with longer reads and algorithms that analyse the raw signal from the sequencing machine.

**Figure 3.**
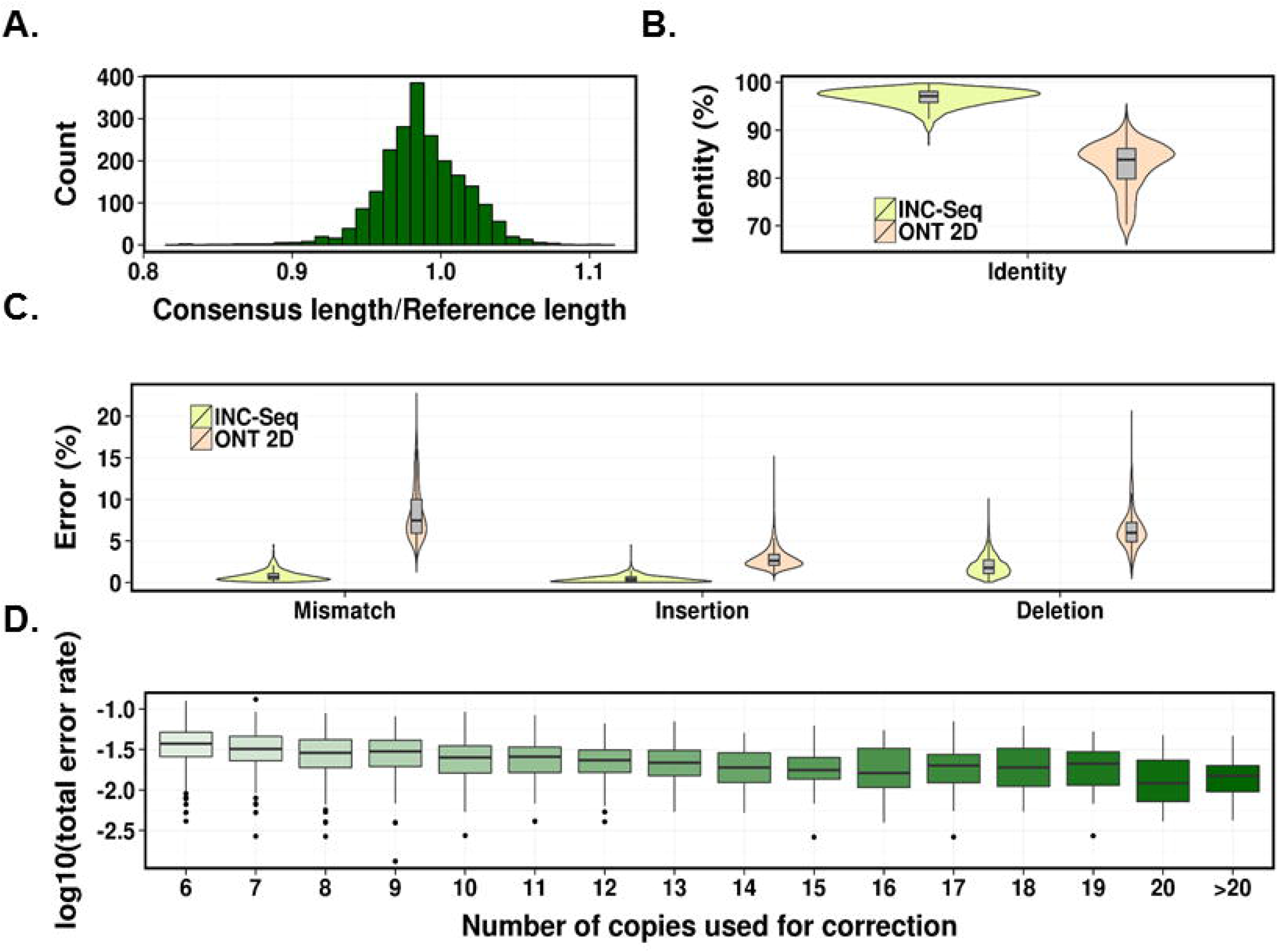
INC-Seq produces long and accurate reads. (A) The ratio between the length of INC-Seq corrected reads and the reference sequence is tightly distributed around one (only reads with length from 600-800bp are shown here). (B and C) INC-Seq boosts overall read accuracy and reduces mismatch, insertion and deletion error rates significantly (one-sided wilcoxon test *p*-value <10^-16^ in all cases). (D) Accuracy of INC-Seq sequences increases with the number of segments used for consensus construction.

### 3.4. MinION INC-Seq reads provide high-accuracy, full-length 16S amplicon sequences

MinION sequencing of the INC-Seq 16S rRNA library provided similar estimates of inter-molecular ligation and template switching rates as was observed for PacBio libraries (**Supplementary Table 1**). Consensus sequences were found to span the 16S amplicons (median and mean coverage: 99%; **Figure 3A**), boosting median accuracy of ONT 2D reads from 84% to 97% (**Figure 3B**). Overall, mismatch rates were reduced 10-fold (from 7.5% to 0.7%; one-sided wilcoxon test *p*-value <10^-16^), with significantly improved insertion and deletion error rates as well (one-sided wilcoxon test *p*-value <10^-16^; **Figure 3C**). Total error rate of consensus reads was found to be negatively correlated with the number of segment copies used to construct the consensus (Spearman *ρ* = -0.37, *p*-value <10^-16^), but plateaued out after 15 copies (**Figure 3D**).

Despite the potential for amplification bias in INC-Seq, relative abundance estimates were found to be surprisingly close to the defined abundances for the simple community on both sequencing platforms (**Table 1**). This could partially be explained by the use of long molecules (containing at least six copies of the amplified template) for INC-Seq analysis, which may be less affected by bias due to exponential amplification. In addition, several long consensus sequences (∼2 kb) were obtained on both sequencing platforms. Many of these were aligned using BLAST to genomic DNA of *streptococci* (92% identity), indicating that INC-Seq can successfully capture longer templates in the input as well.

### 3.5. INC-Seq enables robust and sensitive 16S rRNA profiling

To further evaluate INC-Seq’s utility for 16S rRNA profiling, we applied it to a synthetic community of 10 microbial species with widely varying relative abundances and where some bacteria share high similarity (>95%; **Table 2**, **Figure 4A**). To check reproducibility, two RCA replicates were generated with the same pool of bacterial DNA and consensus INC-Seq reads were mapped to a comprehensive 16S database with BLAST (see **Section 2.6**). Since 16S references can be highly similar, we applied a stringent filter (reference coverage >0.98 and identity >0.98) to remove alignments with low confidence. The resulting abundance profiles were found to be consistent across replicates (**Figure 4B**; Pearson *ρ*=0.99, *p*-value<10^-8^) and thus combined for species profiling. In addition, the merged profile was significantly correlated with defined abundances (Pearson *ρ*=0.83, *p*-value=0.003). Relative abundance estimates for the most abundant species in the community were however significantly off from defined values suggesting that INC-Seq libraries for the staggered community might have been more affected by amplification bias, though experimental error in community composition cannot be ruled out. Interestingly, except for one read that was ambiguously classified because it was mapped to *K. pneumoniae* and *K. variicola* (differ by 1 nucleotide) with the same identity, no other false positives were detected (**Table 2**). Consensus INC-Seq reads could be used to estimate the abundance of *S. epidermidis* to be 0.7% (defined abundance = 0.5%) despite the presence of *S. aureus* in the community (99% identical with defined abundance = 36%; **Figure 4A**). Despite the stringent filters used, INC-Seq could be used to identify all species present in the community, including *F. prausnitzii* with a defined relative abundance of 0.1% (**Table 2**).

**Figure 4.**
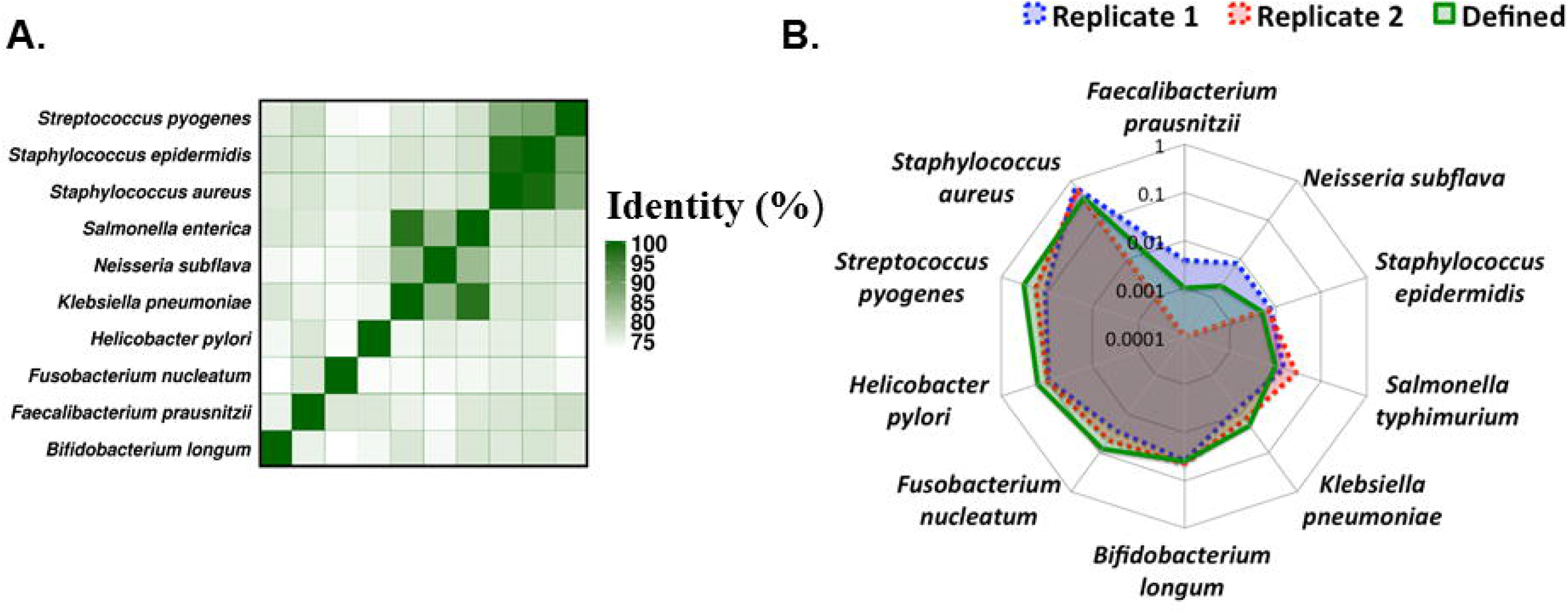
Species detection using INC-Seq. (A) Ten species were selected to construct an artificial community. Some species have highly similar 16S sequences (e.g. *S. aureus* and *S. epidermidis* share 99% identity). (B) Two separate INC-Seq runs produce consistent results that are well correlated with defined abundances (Pearson ρ=0.83, *p*-value=0.003).

## 4. Discussion

The INC-Seq protocol is relatively easy to implement and thus could potentially be incorporated into automated library preparation workflows. For a modest increase in experimental cost (a few hundred dollars), it provides more accurate long reads on the MinION platform, allowing for the use of nanopore sequencing for a wider range of applications. We are currently exploring variations of INC-Seq, including the use of dumbbell adaptors to further refine it and simplify the bioinformatics workflow. Two areas of particular focus are (i) the ability to generate accurate consensus reads for variable-sized fragments without significantly distorting relative abundance estimates and (ii) using longer raw reads to push consensus read accuracy higher.

We explored the use of INC-Seq for 16S rRNA based profiling as it is a widely used approach to investigate complex bacterial communities. Being a targeted approach, it reduces sequencing cost (particularly when non-microbial contamination is high) and is able to capture rare species in a community. Long and accurate INC-Seq reads are well suited to distinguish highly similar 16S sequences allowing for accurate classification at the species level. Amplification of longer regions may improve resolution further, but with potential reduction in the set of sequences that can be amplified [10]. INC-Seq libraries were found to be able to identify rare species in the community but further experimentation would be needed to understand the limits of its sensitivity with greater sequencing depth [18].

Several steps in the analysis of raw INC-Seq reads could be improved further. The identification of segments to construct the consensus was designed to be conservative, but may result in many informative reads and segments being discarded. Segments from chimeric reads could also be used to construct more than one consensus sequence but we conservatively excluded this option. In principle, with the use of six or more segment copies, INC-Seq consensus reads could be more accurate than observed, but this likely requires the development of consensus approaches that realign and analyze sequences at the signal level [19, 20].

Long and accurate reads are key to many genomic applications, e.g. in the case of human haplotype reconstruction where polymorphisms are typically separated by a kilobase in the genome. With long and accurate reads, the computational requirement for phasing is greatly reduced [3]. Other applications include exon connectivity detection [2] and viral quasi-species analysis [21, 22]. With the release of PromethION, a highly scalable and flexible real-time sequencing and analysis platform (300× more pores than the MinION system; https://www.nanoporetech.com/products-services/promethion), we anticipate that nanopore sequencing will be cost-effective for many of these applications, and more practical based on improved accuracy using INC-seq.

---

**Supplementary Figure 1.**
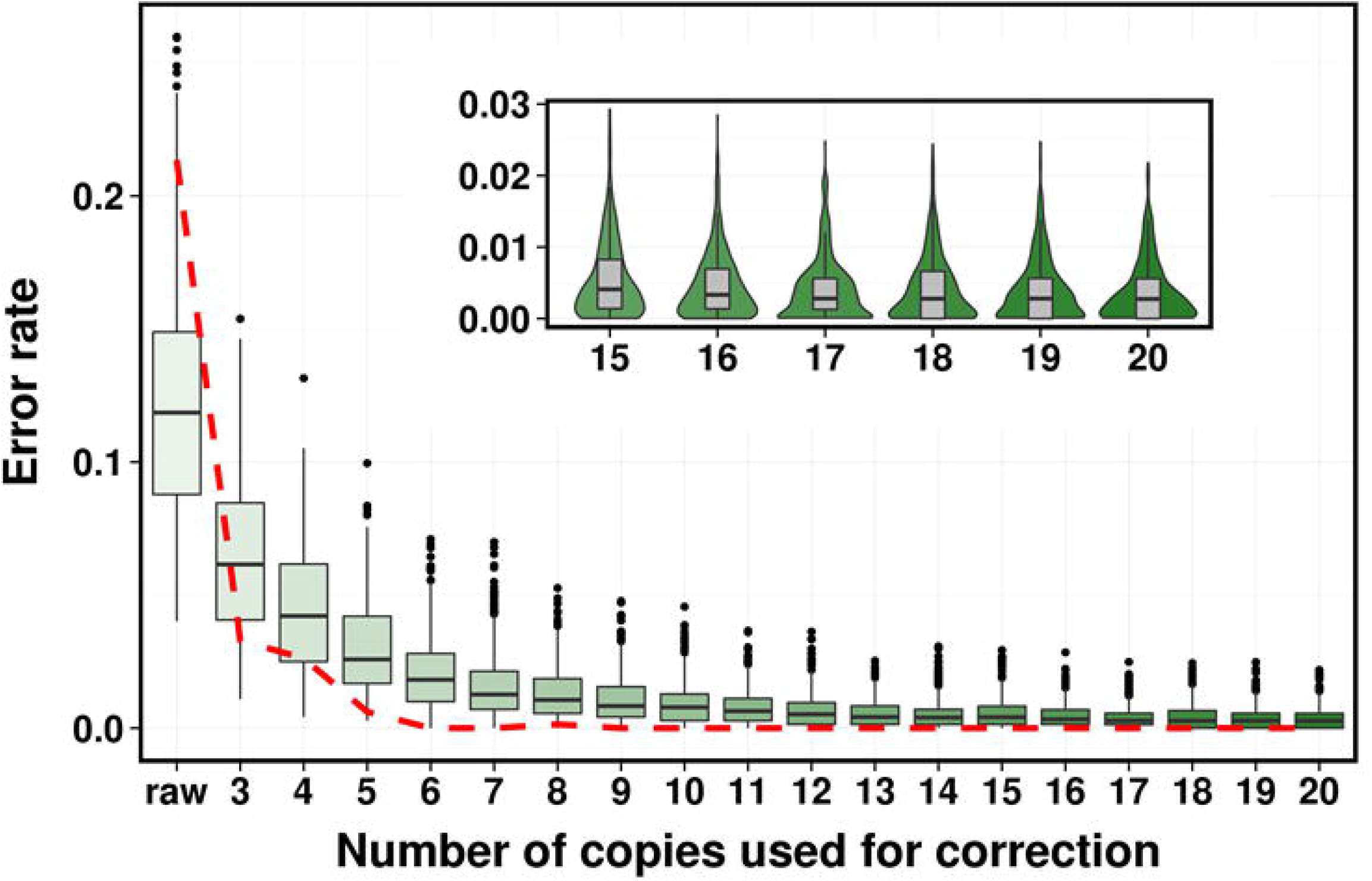
Error rates of simulated raw and INC-Seq corrected reads. Boxplots represents 10 separate simulations from a randomly selected OTU reference sequence. The inner panel is a violinplot for the right tail of the boxplot. The dashed curve shows the change in average OTU assignment accuracy.

**Supplementary Figure 2.**
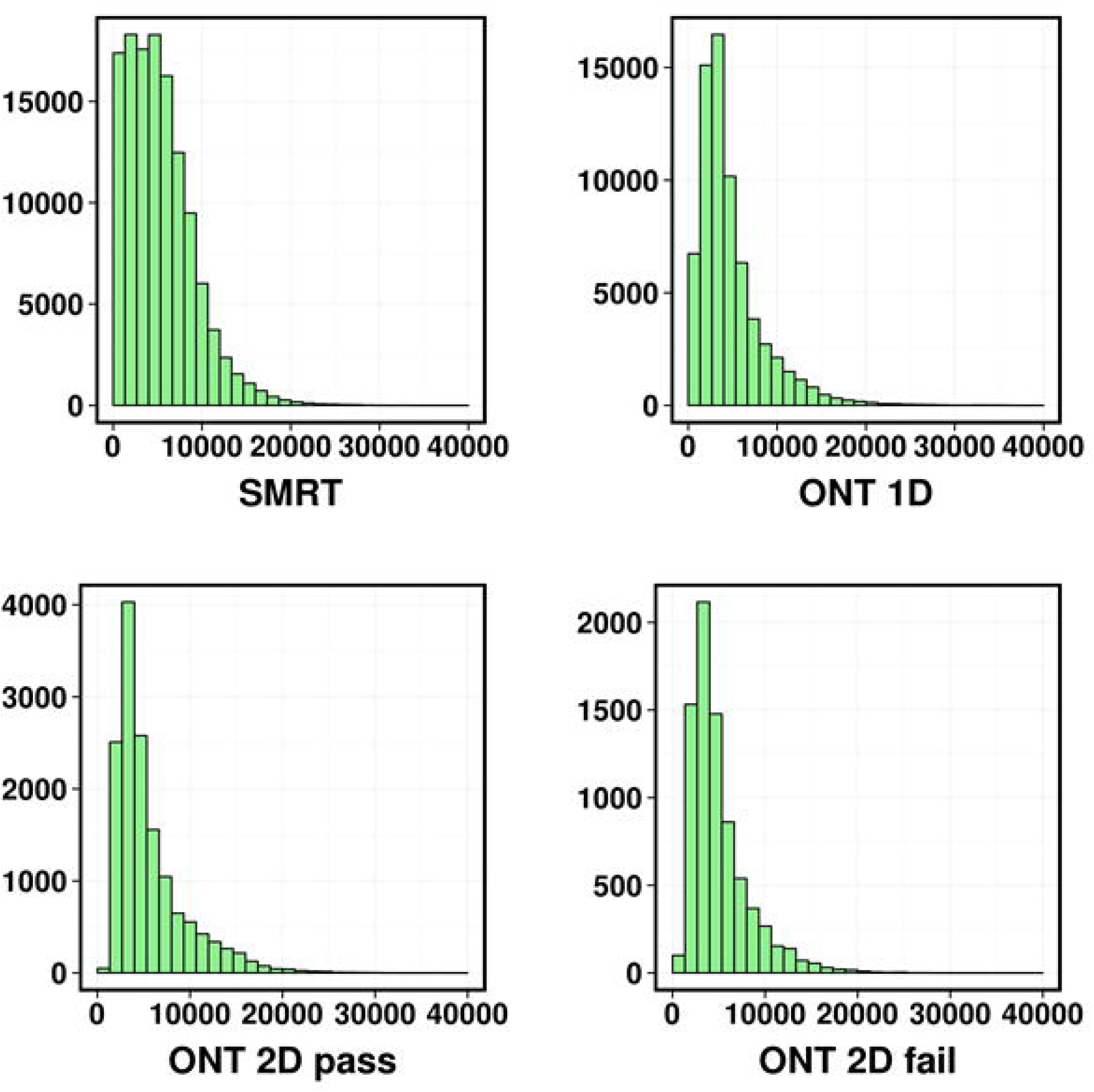
Read length distribution of SMRT and ONT reads (read length on x-axis and counts on y-axis).

**Supplementary Figure 3.**
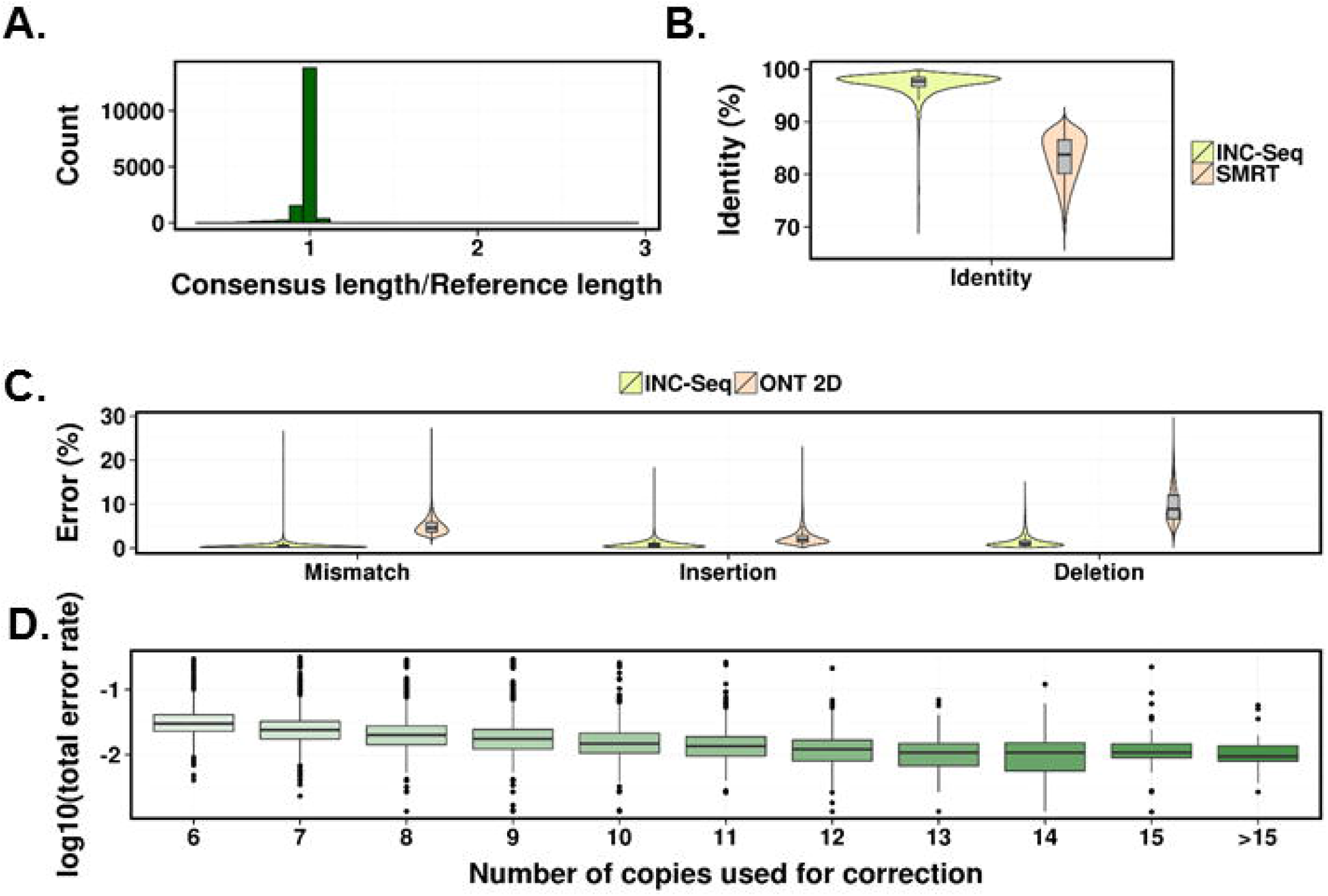
INC-Seq performance on SMRT reads (A) The ratio between the length of INC-Seq corrected reads and reference sequences is tightly distributed around one (mean: 0.982; standard deviation 0.055). (B and C) INC-Seq boosts overall read accuracy and reduces mismatch, insertion and deletion error rates significantly. (D) Accuracy of INC-Seq sequences increases with the number of segments used for consensus construction.

**Supplementary Table 1.** Sequencing statistics and chimera rates

|  | Nanopore<br>(Simple) | Pacbi<br>o | Nanopore (Ladder) |  |
| --- | --- | --- | --- | --- |
|  |  |  | Replicate 1 | Replicate 2 |
| **Total number of reads** | 45268 | 126456 | 27937 | 7989 |
| **Number of 2D pass reads** | 14583 | - | 7444 | 2904 |
| **Estimated template switching rate** | 0.006 | 0.028 | - | - |
| **Estimated inter-molecular ligation rate** | 0.0009 | 0.0006 | - | - |
| **Number of reads longer than 4kb** | 7993 | 73187 | 2909 | 2074 |
| **Number of reads corrected** | 2177 | 16327 | 1076 | 1183 |

